# Tumor Microenvironmental Interactions of Mast Cells and Lymphocytes in Gastric Cancer: A Tissue-Based Study

**DOI:** 10.1101/2025.08.05.668620

**Authors:** Gazi Abdus Sadique, Md. Shahriar Mamun

## Abstract

**Background:** Mast cells and tumor-infiltrating lymphocytes (TILs) are key immune components in the tumor microenvironment (TME). Their densities and interaction dynamics may influence gastric cancer progression. This research intends to investigate the density and prevalence of mast cells and CD8+ tumor-infiltrating lymphocytes in gastric adenocarcinoma while evaluating their correlation with tumor grades.

**Methods:** This cross-sectional observational study was conducted in the Department of Pathology, Satkhira Medical College, from August 2024 to July 2025. Ethical clearance was obtained from the Ethical Review Committee of Satkhira Medical College prior to study initiation. A total of 50 cases of invasive gastric adenocarcinoma were included. Routine Hematoxylin and Eosin (H&E) staining was performed for grading. Mast cell density was assessed using toluidine blue staining and CD8+ TILs by immunohistochemistry. Associations with tumor grades were evaluated using the Chi-square test.

**Result:** Mast cell density significantly increased with higher tumor grades (p<0.05). CD8+ TILs, both intra-and peri-tumoral, also showed a statistically significant synergistic higher distribution across tumor grades (p=0.031 and p=0.012, respectively), with higher TIL infiltration in higher grades.

**Conclusion:** Mast cells and CD8+ lymphocytes are similarly distributed across gastric cancer grades, suggesting their synergistic roles in tumor progression. These findings highlight potential immunoregulatory cross-talk within the TME that may influence therapeutic targeting.

## Background

Stomach cancer continues to be one of the most common and deadly cancers worldwide, especially impacting communities in South Asia, such as Bangladesh. Despite advances in surgical and chemotherapeutic modalities, the overall prognosis for advanced gastric cancer remains poor, largely due to late-stage diagnosis and tumor heterogeneity [1,2]. In recent times, cancer research has shifted more towards the tumor microenvironment (TME) which is a dynamic ecosystem consisting of immune cells, fibroblasts, components of the extracellular matrix (ECM), blood vessels, and a diverse range of cytokines and chemokines that together influence tumor behavior [3].

Among the various immune cells within the TME, mast cells (MCs) and tumor-infiltrating lymphocytes (TILs) have attracted growing attention for their synergistic influence on cancer progression. Mast cells, long recognized for their role in allergy and inflammation, are now understood to be versatile modulators of the tumor microenvironment. They secrete variety of bioactive substances, including tryptase, histamine, vascular endothelial growth factor (VEGF), and interleukins, which can promote angiogenesis, tissue remodeling, and immune evasion [4,5]. Their exact function in cancer, however, remains paradoxical because some studies link high mast cell density to poor prognosis and tumor progression, while others suggest potential antitumor effects, possibly via recruitment of cytotoxic T cells [6].

In contrast, CD8+ cytotoxic T lymphocytes (CTLs) have long been recognized as essential players in the immune response against tumors. They directly eliminate cancer cells by releasing perforin and granzyme, and their presence has been linked to positive clinical results in a range of solid tumors, including gastric cancer [7,8]. Elevated levels of CD8+ tumor-infiltrating lymphocytes (TILs) are associated with lower rates of recurrence and enhanced survival, and are now recognized as a characteristic of an “inflamed” or “hot” tumor, which is likely to respond favorably to immune checkpoint inhibition therapies [9].

Although there is a substantial amount of research on both mast cells and CD8+ T cells separately, their interaction within the gastric tumor microenvironment is still not well understood. New findings from different types of cancer indicate that mast cells might influence lymphocyte infiltration by releasing chemokines (such as CCL5 and CXCL10) or through the release of immunomodulatory cytokines like IL-10 and TGF-β [10,11]. Such interactions could either potentiate or suppress cytotoxic T-cell responses depending on the spatial distribution, activation status, and density of these immune cells. Understanding this interplay is important for understanding mechanisms of immune evasion and may help in identifying potential therapeutic targets.

Therefore, this study aims to explore the density and distribution of mast cells and CD8+ TILs in gastric adenocarcinoma, and to assess their association with tumor grades. This study aims to provide new understanding of the immune structure and behavior in gastric cancer, which could have significant effects on prognosis and tailored immunotherapy.

## Materials and Methods

This cross-sectional observational study was conducted in the Department of Pathology, Satkhira Medical College, from August 2024 to July 2025. A total of 50 cases of histologically confirmed gastric adenocarcinoma were included in the study. Ethical clearance was obtained from the Ethical Review Committee of Satkhira Medical College prior to study initiation. Demographic information, including age and sex, as well as tumor histologic grade, was collected from institutional records and pathology requisition forms. Formalin-fixed, paraffin-embedded (FFPE) tissue blocks were sectioned at 5 µm and stained using routine Hematoxylin and Eosin (H&E) for initial morphological evaluation. To assess mast cell density, sections were stained with toluidine blue. Mast cells were identified by their violet to dark purple cytoplasmic granules and counted under high-power fields (HPFs, 400× magnification). The average number of mast cells per HPF was calculated from five randomly selected fields in each section. Immunohistochemical (IHC) staining for CD8 was performed to identify cytotoxic T lymphocytes. IHC was conducted using a monoclonal anti-CD8 antibody, following standard antigen retrieval and DAB-based detection protocols. CD8+ lymphocytes were quantified separately in the intra-tumoral and peri-tumoral regions by counting five randomly selected HPFs. Infiltration levels were categorized into low and high based on median values across the cohort. All statistical analyses were performed using SPSS software (version 26). Descriptive statistics were used for demographic and histopathological variables. The Pearson Chi-square test was employed to evaluate the associations between mast cell density, CD8+ lymphocyte infiltration, and tumor grade. A p-value < 0.05 was considered statistically significant.

## Results and observation

This study included 50 histologically confirmed cases of gastric adenocarcinoma. The mean age of the patients was 56.89 ± 9.55 years (range: 30–75 years), with the majority (58%) falling within the 40–60-year age group. Male predominance was noted, with 56% male and 44% female patients. Histologically, the tumors were most frequently classified as Grade 2 (60%), followed by Grade 3 (26%) and Grade 1 (14%). Mast cell density was significantly associated with tumor grade (*p* < 0.05), showing a progressive increase from Grade 1 (mean: 3.83 ± 1.18) to Grade 3 tumors (mean: 11.25 ± 1.24). Analysis of CD8+ tumor-infiltrating lymphocytes (TILs) revealed that intra-tumoral CD8+ TILs were high in 66% of cases, with a statistically significant synergistic relationship with tumor grade (*p* = 0.031); high intra-tumoral CD8+ TILs were predominantly seen in Grade 3 tumors. Similarly, peri-tumoral CD8+ TILs were elevated in 70% of cases, with a significant association between higher TIL density and higher tumor grade (*p* = 0.012). Grade 1 tumors showed the lowest frequency of high peri-tumoral CD8+ TILs. These findings suggest that both mast cell density and CD8+ TIL distribution are closely related to the histopathological grading of gastric adenocarcinoma.

**Table 01:** Distribution of the study subjects by their age (n = 50).

| Age (years) | Frequency | (%) |
| --- | --- | --- |
| <40 | 4 | 8.0% |
| 40-60 | 29 | 58.0% |
| >60 | 17 | 34.0% |
| Mean ± SD | 56.98 ± 9.55 | |
| Range (Min-Max) | (30-75) |  |

**Figure 01:**
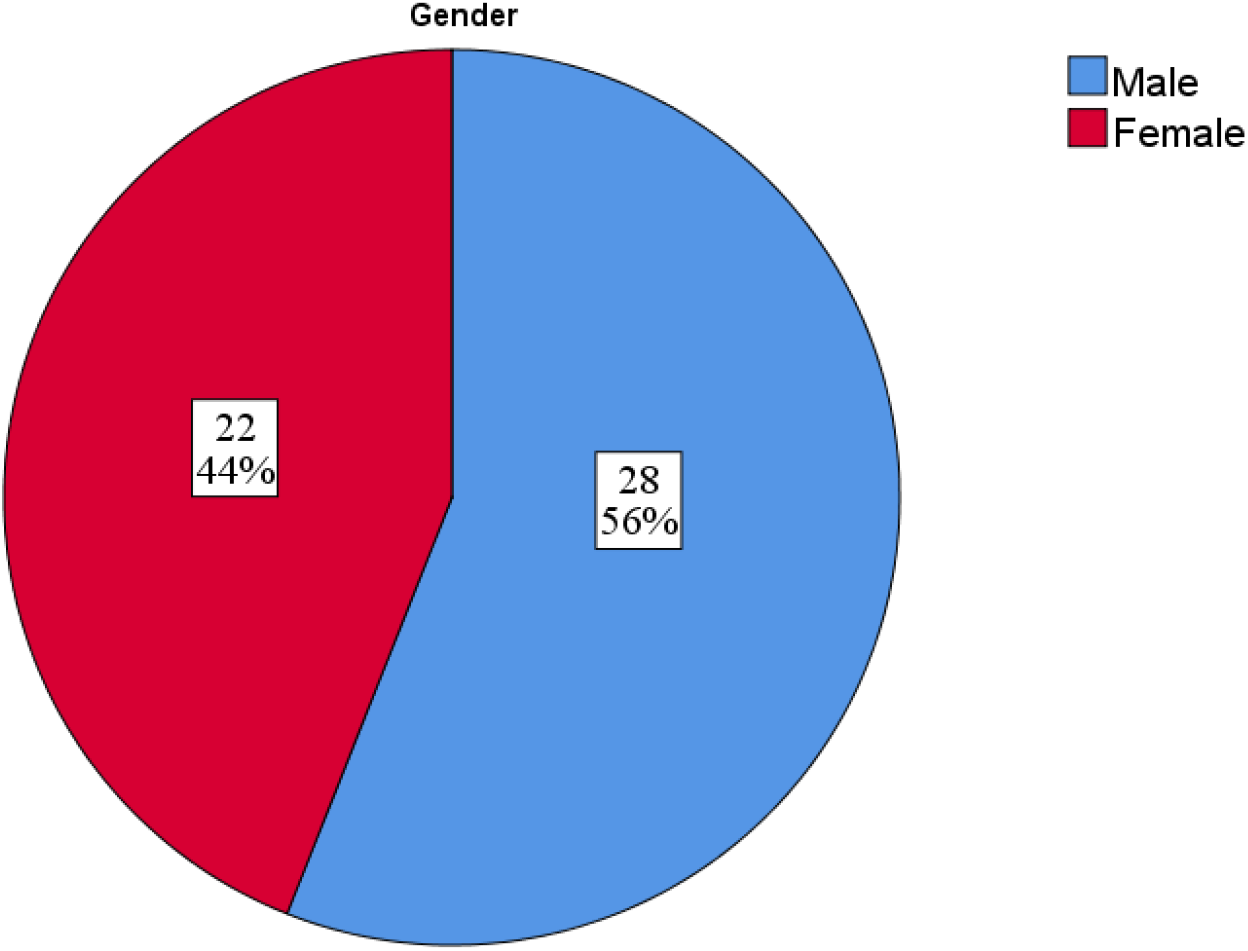
Distribution of cases according to gender (n=50)

**Table 02:** Distribution of the study subjects by tumor grade (n = 50).

| Grades | Frequency | Percent |
| --- | --- | --- |
| Grade 1 | 7 | 14.0 |
| Grade 2 | 30 | 60.0 |
| Grade 3 | 13 | 26.0 |
| **Total** | 50 | 100.0 |

**Table 03:** Mast cell density among the study groups (n=50).

| **Mast cell density** | **Frequency** | **Percent** |
| --- | --- | --- |
| Low mast cell density | 25 | 34.0 |
| High mast cell density | 25 | 66.0 |
| **Mean ± SD** | **7.13 ± 2.85** |  |
| **Median** | **6.66** |  |
| **Range (Min-Max)** | **(2.00-12.66)** |  |

**Table 04:** Association between mast cell density and grade of tumor (n=50).

| **Grades** | **Mast cell density** | **p-value** |
| --- | --- | --- |
| Grade 1 | 3.85 ± 1.18 | <0.05 |
| Grade 2 | 6.11 ± 1.24 | |
| Grade 3 | 11.25 ± 1.24 | |

**Table 05:**
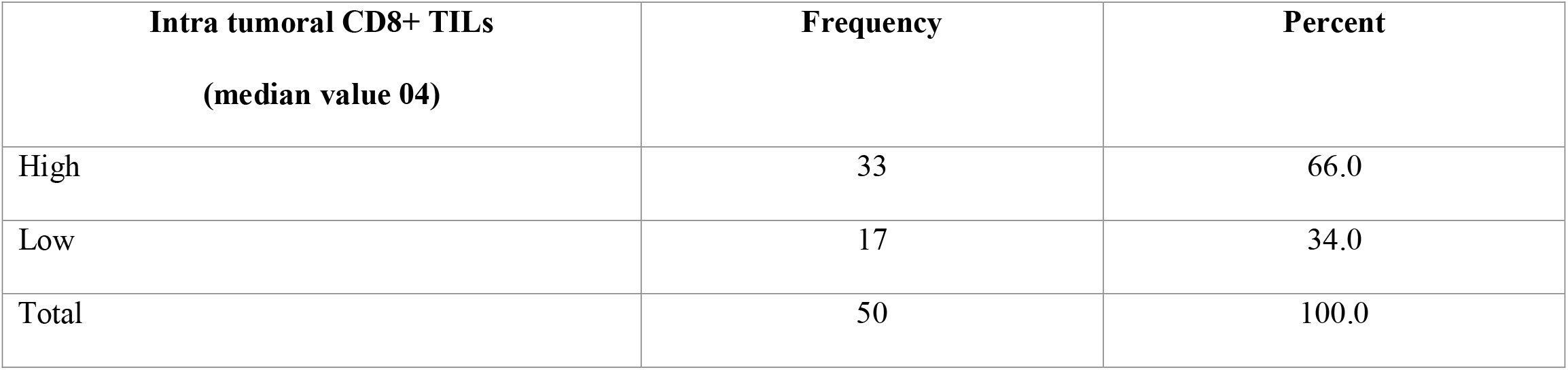
Distribution of the cases according to CD8+ intra tumoral TILs (n=50).

**Table 06:**
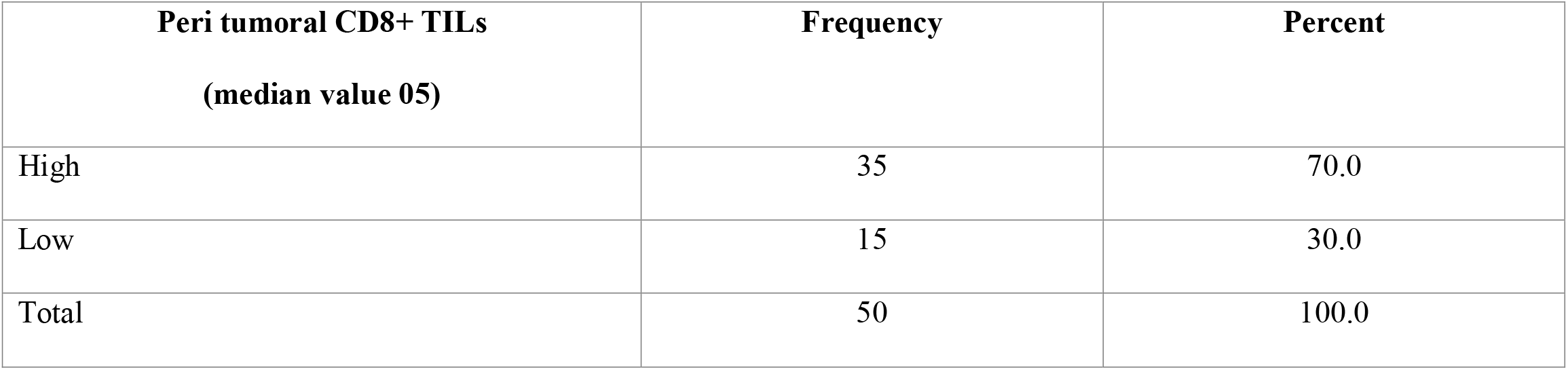
Distribution of the cases according to CD8+ peri tumoral TILs (n=50).

**Table 07:**
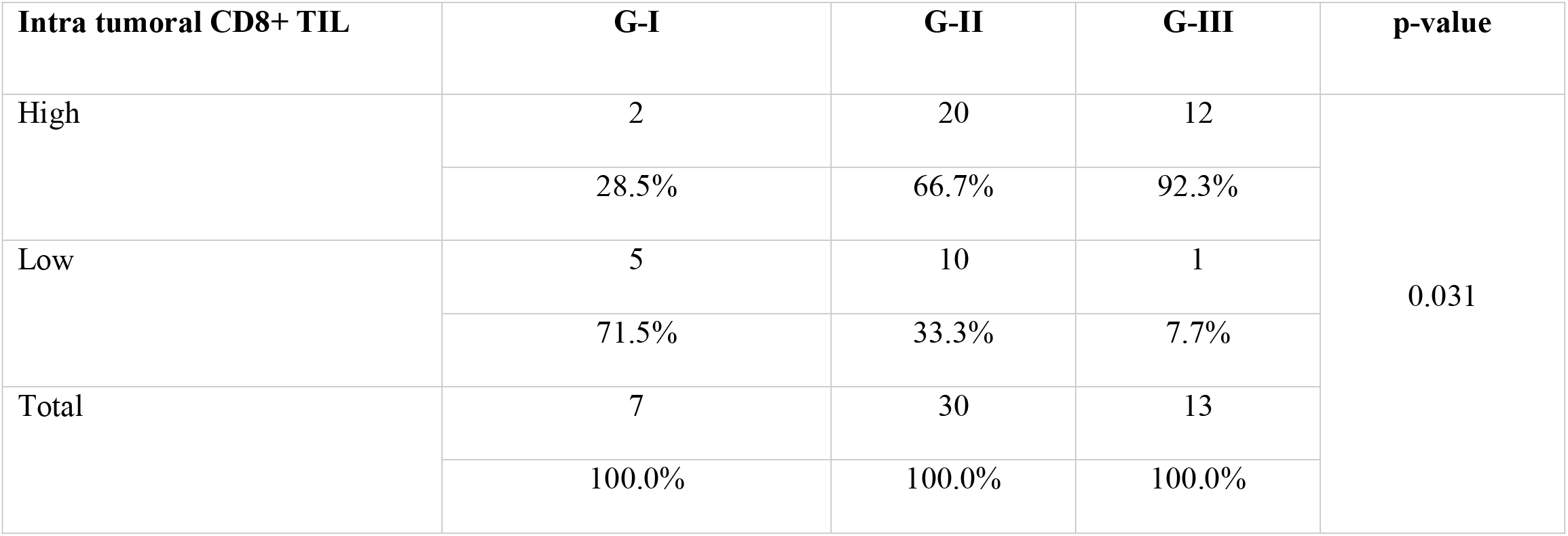
Association of intra tumoral CD8+ TILs with histopathological grades of gastric adenocarcinoma (n=50).

**Table 08:** Association of peri tumoral CD8+ TILs with histopathological grades of gastric adenocarcinoma (n=50).

| **Peri tumoral CD8+ TIL** | **G-I** | **G-II** | **G-III** | **p-value** |
| --- | --- | --- | --- | --- |
| High | 1 | 19 | 13 | 0.012 |
|  | 14.3% | 63.3% | 100% |  |
| Low | 6 | 11 | 0 |  |
|  | 85.7% | 36.7% | 0% |  |
| Total | 7 | 30 | 13 |  |
|  | 100.0% | 100.0% | 100.0% |  |

**Figure 02:**
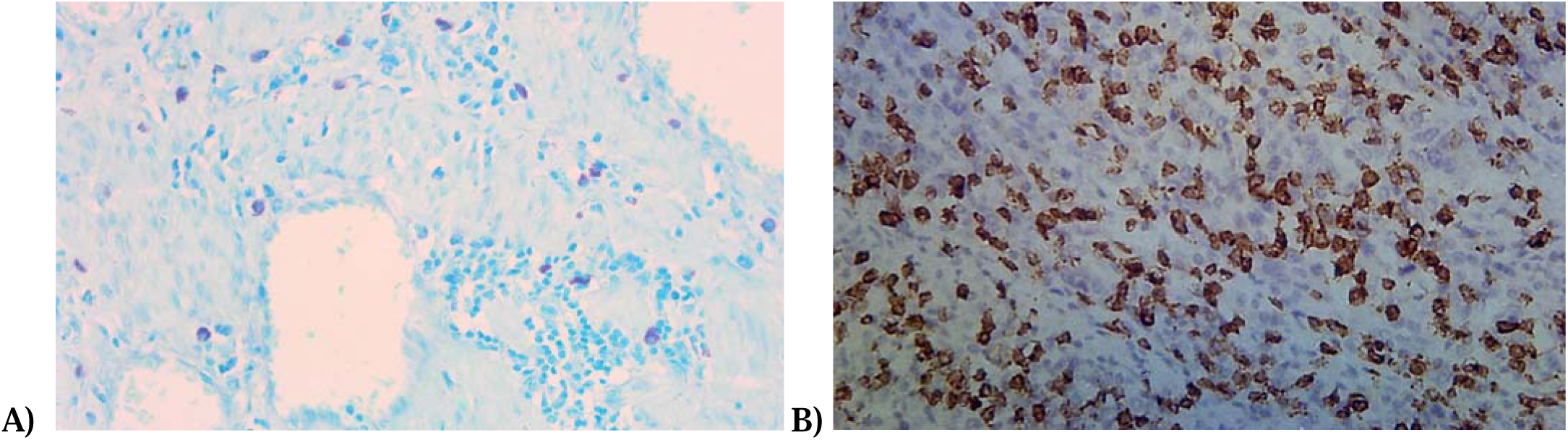
A. High infiltration of mast cells in grade 3 gastric adenocarcinoma B. High CD8 IT-TIL infiltration in grade 3 gastric adenocarcinoma.

## Discussion

The present study aimed to investigate the association of mast cell density and CD8+ tumor-infiltrating lymphocytes (TILs), both intra- and peri-tumoral, with the histopathological grade of gastric adenocarcinoma. Our findings demonstrate a significant correlation between immune cell infiltration and tumor grade, reflecting the role of immune modulation in gastric cancer progression.

We observed a male predominance and a peak incidence in the 40–60-year age group, consistent with epidemiological patterns reported in previous studies from South Asia and other developing regions [12,13]. The prevalence of moderately differentiated tumors (Grade 2) corresponds with worldwide patterns, though local dietary habits and exposure to Helicobacter pylori could affect this distribution [14].

Mast cell density was significantly elevated in high-grade tumors (Grade 3), suggesting its potential role in tumor progression. This finding supports existing literature indicating that mast cells, through the release of angiogenic and immunomodulatory factors such as VEGF, TNF-α, and IL-6, may contribute to tumor growth, immune evasion, and extracellular matrix remodeling [15, 16]. Although mast cells have typically been viewed as components of the host defense system, they seem to take on a pro-tumorigenic function within the tumor microenvironment, especially in advanced cancers [17].

Interestingly, CD8+ TIL density was also found to be higher in poorly differentiated tumors, contrasting with conventional views that associate high T-cell infiltration with early-stage, well-differentiated tumors and improved prognosis [18]. This paradoxical finding may indicate that higher-grade tumors, due to their high mutational burden and neoantigen load, stimulate a more intense immune response [19]. Spatial analysis further revealed increased densities of both intra- and peri-tumoral CD8+ TILs in high-grade tumors. While intra-tumoral CD8+ TILs are typically linked with active tumor cell targeting, peri-tumoral accumulation may reflect an immunological bottleneck or exclusion phenomenon, where effector T cells are recruited but fail to penetrate the tumor core [20]. These findings resonate with prior observations in melanoma and colorectal cancer, where TIL location correlates with immune escape mechanisms and therapeutic response [21].

Previous studies have demonstrated both favorable and unfavorable prognostic implications of CD8+ TILs in gastric cancer, often depending on their spatial distribution, activation status, and interaction with regulatory immune cells or checkpoints [22]. Therefore, the increase in TILs in high-grade tumors should not be interpreted in isolation as an anti-tumor signal; rather, it may reflect a complex immunological phenomenon between host defense and immune suppression.

Our study benefits from histological quantification of immune infiltrates and correlational analysis with tumor grade. However, limitations include a modest sample size, absence of survival follow-up, and lack of molecular or functional characterization of immune cells. Further studies incorporating immune phenotyping and checkpoint expression profiling are warranted to unveil the biological behavior and clinical implications of these findings.

In conclusion, both mast cell density and CD8+ TILs were positively associated with higher histological grade in gastric adenocarcinoma, suggesting their involvement in the progression of more aggressive tumor phenotypes. These results question the oversimplified classification of immune cells as merely pro- or anti-tumor forces and emphasize the necessity for detailed immunological profiling in gastric cancer for both evaluating prognosis and determining therapeutic strategies.

## Conclusion

This study demonstrates a significant association between higher histological grade of gastric adenocarcinoma and increased mast cell density as well as elevated intra- and peri-tumoral CD8+ TIL infiltration. While mast cells appear to support tumor progression through pro-inflammatory and angiogenic mechanisms, the presence of increased CD8+ TILs in poorly differentiated tumors may reflect an ineffective or dysregulated immune response rather than effective tumor control. These findings challenge conventional assumptions about immune cell infiltration as solely protective and signify the complexity of the tumor-immune interplay in gastric cancer. Both mast cell and CD8+ TIL profiles may serve as valuable biomarkers for tumor aggressiveness and could have prognostic and therapeutic implications, particularly in the context of emerging immunotherapeutic strategies. Further studies incorporating functional immune profiling and long-term clinical outcomes are warranted to validate and expand upon these observations.

